# Exploitation of local and global information in predictive processing

**DOI:** 10.1101/687673

**Authors:** Daniel S. Kluger, Nico Broers, Marlen A. Roehe, Moritz F. Wurm, Niko A. Busch, Ricarda I. Schubotz

## Abstract

While prediction errors have been established to instigate learning through *model adaptation*, recent studies have stressed the role of model-compliant events in predictive processing. Specifically, so-called *checkpoints* have been suggested to be sampled for *model evaluation*, particularly in uncertain contexts.

Using electroencephalography (EEG), the present study aimed to investigate the interplay of such global information and local *adjustment cues* prompting on-line adjustments of expectations. Within a stream of single digits, participants were to detect ordered sequences (i.e., 3-4-5-6-7) that had a regular length of five digits and were occasionally extended to seven digits. Across experimental blocks, these extensions were either rare (low *irreducible uncertainty*) or frequent (high uncertainty) and could be unexpected or indicated by incidental colour cues.

Exploitation of local cue information was reflected in significant decoding of cues vs non-informative analogues using multivariate pattern classification. Modulation of checkpoint processing as a function of global uncertainty was likewise reflected in significant decoding of high vs low uncertainty checkpoints. In line with previous results, both analyses comprised the P3b time frame as an index of excess model-compliant information sampled from probabilistic events.

Accounting for cue information, an N400 component was revealed as the correlate of locally unexpected (vs expected) outcomes, reflecting effortful integration of incongruous information. Finally, we compared the fit of a *global model* (disregarding local adjustments) and a *local model* (including local adjustments) using representational similarity analysis (RSA). RSA revealed a better fit for the global model, underscoring the precedence of global reference frames in hierarchical predictive processing.

## Introduction

One of the central faculties of the human brain is to predict upcoming events based on internal models of the world. Incoming sensory information is constantly compared to model-based predictions and resulting mismatches - termed *prediction errors* (PE) - are used to continuously update the model for future reference (Friston, 2005; Mumford, 1992). Consequently, as such unexpected events are particularly informative, the importance of PE for associative learning through model adaptation has long been established (Bastos et al., 2012; Den Ouden et al., 2009). Recent studies have stressed the role of probabilistic events that carry behaviourally relevant information but do not indicate a mismatch (Kühn & Schubotz, 2012; Kluger et al., 2019; Trempler et al., 2017). In contrast to *model adaptation* instigated by PE, these critical points in time have been suggested to serve outcome-independent *model evaluation*. On-line exploitation of model-compliant sensory input at so-called *checkpoints* (CP) has been shown to be modulated by higher-order context statistics such as irreducible uncertainty with regard to outcome expectancies (Kluger & Schubotz, 2017): In a highly uncertain environment, additional resources are allocated to the processing of informative events to resolve ambiguity regarding the validity of the internal model – even if these events are model-compliant. Notably, this effect was not merely driven by event-related surprise but instead reflected exploitation of excess information from probabilistic non-error events (i.e., checkpoints).

A subsequent EEG study (Kluger et al., 2019) further demonstrated electrophysiological commonalities and distinctions between congruous and incongruous information processing. When compared to fully predictable standard trials, both CP and PE showed a significant P3b component, marking the joint (im)probabilistic - and therefore, informative - nature of both event types in accordance with the component’s functional significance (Mars et al., 2008; Seer et al., 2016). The direct comparison of checkpoints and prediction errors revealed a significant N400 component indicating a mismatch signal exclusively induced by PEs. In sum, these findings illustrate two important aspects of predictive processing: First, informative model-compliant events appear to take on a similar, but not identical role as canonical mismatch signals in resolving ambiguity with regard to model validity. Second, the extent to which such model-compliant events are exploited to evaluate the internal model is modulated by global, higher-order parameters such as environmental uncertainty. These findings indicate that non-error information has the potential to play a central role in predictive processing and is thus worth a more thorough determination, especially in relation to mismatch signals.

Extending the findings reported so far, the present EEG study aimed to address the potential influence of learned *local* contingencies on predictive processing. Normally, instructive events such as checkpoints and prediction errors are informative due to their probabilistic nature, i.e. their occurrence is unknown before it is actually observed. However, these critical events can conceivably be rendered uninformative by local cues reliably indicating a certain outcome - much like a priority road sign telling drivers not to slow down at a particular intersection. We employed a modification of the paradigm from Kluger et al. (2019) to assess both the electrophysiological signature of these local cues as well as their influence on the processing of expected and unexpected outcomes.

Trained participants performed a serial pattern detection task in which they were to press and hold a response button whenever they detected an ordered digit sequence (e.g. 3-4-5-6-7) within an otherwise pseudorandom stream of coloured single digits. Ordered sequences had an expectable length of five digits but were occasionally extended to be seven digits long. This way, the sixth position could either contain a random digit denoting the regular (i.e., expected) end of the sequence (REG) or an unexpected sequential digit indicating an extension (EXT; see Fig. 1). Relative presentation rates of REG and EXT were probabilistically modulated by blockwise manipulation of *irreducible uncertainty*. This particular measure quantifies uncertainty that cannot be reduced further and remains even after successful (i.e., ideal) learning (De Berker et al., 2016; Payzan-LeNestour & Bossaerts, 2011). Within the scope of the present study, high uncertainty blocks contained an equal number of regular and extended sequences, thus maximising uncertainty with regard to the continuation of a given sequence (Fig. 1B). Crucially, however, some sequences contained colour-coded *adjustment cues* (AC) at the third position. These cues (AC+) provided excess information regarding the continuation of the sequence, reliably indicating a sequential extension (*p* = .75) and thus effectively overriding expectations based on the global task structure.

**Figure 1.**
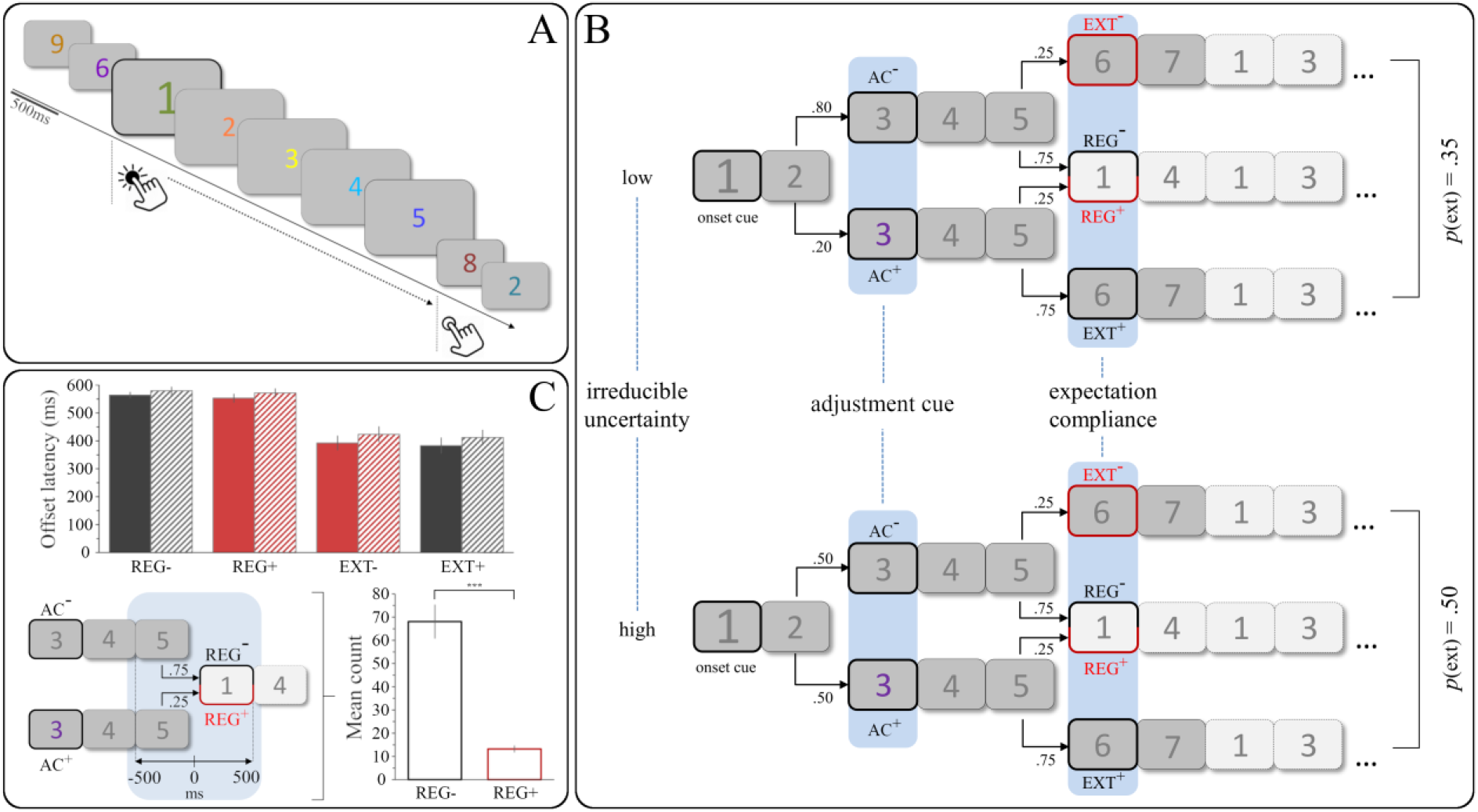
(A) Exemplary trial succession and time frame of the corresponding response for ordered sequences. (B) Experimental manipulations and resulting transition probabilities between trials. Bold framing indicates events of interest for EEG analyses; red framing indicates relatively unexpected events. (C) Top: Mean offset latencies for regular and extended sequences as a function of adjustment cueing and irreducible uncertainty (solid bars = low, hatched bars = high). For significance of main effects, please see main text. Bottom left: Schematic of sampling premature and quick button releases for uncued (REG−) and cued regular sequences (REG+). Bottom right: Mean count of releases within [−500, 500] ms for REG− and REG+ sequences. Error bars show standard error of the mean (SEM). AC = adjustment cue, REG = regular, EXT = extended, *** = p < .001.

Using a similar experimental paradigm without adjustment cues, our previous EEG study (Kluger et al., 2019) had revealed a significant N400 component for the direct comparison of PE and CP. In other words, events that violated cue-based expectations (PE) distinctly elicited an N400 when compared to informative events containing model-compliant sensory information (CP). Since the present study comprised learned adjustment cues (AC) incidentally prompting an on-line shift in model-based expectations, both REG and EXT events could now be locally unexpected as a function of preceding cue information: As AC reliably indicated sequential extensions (EXT+), regular endings following a cue (REG+) were rendered unexpected (see Fig. 1B). Conversely, in sequences without a cue, regular endings (REG−) were the locally expected outcome and extensions (EXT−) were unexpected. Thus, we aimed to replicate previous findings from the direct contrast of expected and unexpected sequence endings within the N400 time frame (Kluger et al., 2019).

Furthermore, we used multivariate pattern classification of event-related potentials (ERP) to conduct two analyses on local vs global modulations: First, one aim of the present study was to assess functional characteristics of the newly introduced adjustment cues. Conceivably, when compared to non-informative third position trials (AC−), predictive information provided by adjustment cues (AC+) could manifest in an electrophysiological signature similar to the one reported by Kluger et al. (2019) for both congruent and incongruent informative events. As our previous study had revealed a joint P3b component to reflect the informativity of both CP and PE, we hypothesised the classifier to reliably decode AC+ and AC− trials within the P3b time frame.

Second, we aimed to validate global effects of checkpoint processing as a function of high vs low uncertainty from our initial fMRI study (Kluger & Schubotz, 2017). Consequently, we sampled regular endings of sequences that did not contain an adjustment cue (REG−), as these events were the precise equivalent of CP from earlier studies. Consolidating previous findings from fMRI and EEG, we hypothesised significant classifier accuracy of *CP high vs CP low* within the P3b time frame (300-500 ms).

Finally, we applied representational similarity analysis (RSA; Kriegeskorte et al., 2008) to test the explanatory power of competing theoretical models relating the representational structures of critical events (AC, CP, PE) to one another. With that goal in mind, we devised two competing binary model representational dissimilarity matrices (RDMs) to predict the empirical representational similarity of event classes in the EEG data. A *global model* categorically assumed high similarity within and high dissimilarity between the sampled event classes (AC+/− vs REG+/− vs EXT+/−) and thus represented global statistics acquired by means of statistical learning (“Sequences are usually five digits long”). In contrast, a *local model* was set up to reflect these adjustments in expectations due to local information: For sequences containing an adjustment cue (AC+), extensions were no longer unexpected (as in the global model) but rather the expected outcome. In these cases, regular sequence endings (REG+) – otherwise the most probable outcome of a sequence – violated the most recent set of expectations based on the adjustment cue (see above). Consequently, we assumed high similarity of events that violated these potentially adjusted predictions following an adjustment cue (REG+, EXT−) in contrast to those that complied with the adjustment cue (REG−, EXT+).

## Materials and methods

### Participants

A total of 30 neurologically healthy, right-handed volunteers (25 female) at the age of 24.8 ± 3.46 years (M ± SD) participated in the study for payment or course credit. Participants were recruited from the university’s volunteer database and had (corrected-to-) normal vision. Written informed consent was obtained from all participants prior to the start of experimental procedures. Experimental standards complied with the local Ethics Committee of the University of Münster.

### Stimulus material

Participants watched pseudorandomly coloured single digits (0 – 9) presented for 500 ms in the centre of a light grey computer screen. Presentation frequencies for all colours and digits were equally distributed both within and across experimental blocks of approximately six minutes. Each block contained *ordered sequences* in which the previous digit was continually increased by one (Fig. 1A). Ordered sequences had a *regular* length of five digits and were embedded in series of *random trials* with no discernible relation between consecutive digits. In order to balance sequential starting points across digits, the ascending regularity necessarily included the 0 character and continued in a circular fashion after the figure 9 (e.g. 8 – 9 – 0 – 1 – 2).

The first digit of every ordered sequence was displayed at 150% of the usual font size, serving as an explicitly instructed *onset cue*. Importantly, digit colours at the third position of each sequence were used to vary expectations regarding its continuation. Each participant was unknowingly assigned one fixed colour that served as an implicit *adjustment cue* (AC): Presentation of the third sequential digit in AC colour (*AC*+ trials) reliably indicated an *extension* of the respective sequence by two digits (*p*_EXT|AC+_ = .75, *p*_REG|AC+_ = .25). Conversely, when the third sequential trial was presented in any colour but the AC colour (*AC*-trials), extensions were rare (*p*_EXT|AC−_ = .25, *p*_REG|AC−_ = .75; see Fig. 1B). AC colours did neither occur at any other sequential position nor during random trials.

Finally, as outlined above, the composition of regular and extended sequences within a particular block was varied across blocks. This way, the *irreducible uncertainty* of a block was set to be either *low* (with *p*_EXT_ = .35) or *high* (with *p*_EXT_ = .50; see Fig. 1B). Low uncertainty blocks could therefore be seen as statistically stable regarding the expected sequence length whereas highly uncertain blocks provided a less stable statistical context. To ensure constant AC validity, high uncertainty blocks necessarily contained a higher percentage of adjustment cues than low uncertainty blocks.

The experiment was programmed and run using the Presentation 18.1 software (Neurobehavioral Systems, San Francisco, CA, USA).

### Task

Participants were instructed to press and hold the left button of a response box with their right index finger whenever an onset cue marked the beginning of any ordered sequence. At the end of any ordered sequence, participants were asked to release the response button as quickly as possible.

### Experimental procedures

The study was conducted on two consecutive days. On the first day, participants completed a training session to provide them with implicit knowledge of the adjustment cues and the underlying statistical structure of the experiment. The training consisted of one high and one low uncertainty block with a total duration of approximately 13 minutes (including a one-minute break). On the second day, participants completed the EEG session which consisted of eight blocks (four blocks of each uncertainty level) with a total duration of approximately 56 minutes (including a one-minute break after each block). Participants were comfortably sitting on a chair in a darkened, sound-dampened and electrically shielded EEG booth. Experimental procedure and task during the EEG session were otherwise identical to the training session.

### Behavioural data analysis

Statistical analyses of behavioural responses were performed in R (R Foundation for Statistical Computing, Vienna, Austria). First, correct and incorrect responses were aggregated separately for each participant. Incorrect responses were further divided into misses (no response over the course of a sequence) and false alarms (response occurring without presentation of sequential trials). Participants’ overall performances were assessed via the discrimination index P_R_ (probability of recognition; Snodgrass & Corwin, 1988).

Offset latency was calculated as reaction time relative to the onset of the first random trial after a particular sequence. Responses occurring either before the go-cue or more than 1500 ms after the end of the sequence were excluded. Repeated-measures analyses of variance (ANOVA) and paired *t*-tests were used to assess potential differences in offset latencies as a function of expectation compliance, adjustment cues, and irreducible uncertainty.

### EEG data analysis

#### EEG data acquisition and data preprocessing

Scalp EEG was recorded from 62 active Ag/AgCl-electrodes mounted in an actiCAP snap electrode cap (Brain Products, Gilching, Germany) using the BrainVision Recorder software (Brain Products, Gilching, Germany). All scalp channels were measured against a ground electrode at position FPz and referenced against FCz during recording. Two additional electrooculogram (EOG) electrodes were applied next to and below the right eye for the detection of horizontal and vertical eye movements, respectively. EEG was recorded at a sampling rate of 1 kHz with recording filters set to 0.1 – 1000 Hz bandpass. EEG preprocessing was conducted in EEGLAB (Delorme & Makeig, 2004). Data segments containing experimental breaks were discarded prior to independent component analysis (ICA). Resulting components distinctly reflecting eye movements were subsequently rejected manually (mean = 1.8 components) using the SASICA toolbox (Chaumon et al., 2015). Data were then filtered with a 0.1 Hz low cut and 30 Hz high cut filter and recalculated to common average reference. A time frame of [-200, 800] ms was defined for the analysis of event-related potentials (ERP). Epochs containing artefacts were discarded by semiautomatic inspection with an allowed maximum/minimum amplitude of ± 200 μV and voltage steps no higher than 50 μV per sampling. Channels with a high proportion of outliers (kurtosis criterion: z > 6) were replaced by a linear interpolation of their neighbour electrodes (mean = 2.1 interpolated channels).

#### Event-related potentials

Averages of the epochs representing events of interest were calculated separately per participant. The third trial of every ordered sequence was sampled to compare trials with and without the learned adjustment cues (AC+ vs AC−). Since ordered sequences had a regular length of five digits, the subsequent digit was critical with regard to expectation compliance: For regular sequences, this position contained the first random digit and marked the end of the ordered sequence (REG). For extended sequences, this position contained the sixth sequential digit and marked the extension of the ordered sequence (EXT). Both outcomes were further subdivided to reflect whether an adjustment cue had been presented at the third position (REG+, EXT+) or not (REG−, EXT−). Grand average ERPs across participants were calculated for all events of interest.

For stringent hypothesis testing, ERP analyses were restricted to specific time frames. Using the Mass Univariate ERP Toolbox (Groppe et al., 2011), ERPs from the respective conditions were submitted to a repeated measures cluster mass permutation test (Bullmore et al., 1999; family-wise significance level α = .05). Repeated-measures *t*-tests were performed for each comparison using the original data and 5000 random within-participant permutations of the data. For each permutation, all *t*-scores corresponding to uncorrected *p*-values of *p* = .05 or less were then bound into clusters. The sum of the *t*-scores in each cluster defined the “mass” of that cluster and the most extreme cluster mass in each of the 5001 sets of tests was used to estimate the distribution of the null hypothesis.

Attempting to replicate previous N400 findings for expectation violations, we directly compared locally unexpected (EXT−, REG+) and expected outcomes (EXT+, REG−) within the N400 time frame (*UNEXP vs EXP*, 300 – 500 ms).

#### Multivariate classification analyses

Multivariate classification was carried out in Matlab using the CoSMoMVPA toolbox (Oosterhof et al., 2016). To increase signal-to-noise ratio (SNR), EEG time series were first downsampled to 100 Hz (see Carlson et al., 2013). SNR and classifier performance were further increased by continuously averaging *n* = 4 trials from the same respective categories before decoding (Grootswagers et al., 2017; Isik et al., 2013). Classification was then performed separately for each 10 ms bin by means of a temporal searchlight analysis approach based on data of all EEG channels (EOGs excluded). Specifically, a linear discriminant analysis (LDA) classifier was trained and tested on trials from two categories (e.g., AC+ and AC− trials) using a leave-two-trials-out cross-validation scheme. This way, classification analyses yielded time courses of decoding accuracy with 10 ms temporal resolution within the interval of [−200, 800] ms. Threshold-free cluster enhancement (TFCE; Smith & Nichols, 2009) with default parameters was used to determine periods of significant decoding accuracy (with a fixed chance level of *p* = .50 for pairwise decoding). As implemented in CoSMoMVPA, permutation testing with 10,000 null iterations and subsequent thresholding at *Z* > 1.96 (*p* < .05) was used to correct for multiple comparisons.

With regard to functional event signatures, two comparisons were the focus of pairwise decoding analyses. First, we assessed the classifier’s performance decoding AC+ from AC− trials over the ERP time course (see Fig. 1B). Third-position trials containing an adjustment cue (AC+) provided excess information with regard to the continuation of the sequence, effectively overriding expectations based on the global task structure. Exploitation of this particular cueing information should translate to distinct electrophysiological correlates when compared to non-cue trials at the same position (AC−). As we had previously reported P3b effects for the comparison of model-compliant, particularly informative events with non-informative equivalents (Kluger et al., 2019), we hypothesised significant above chance classifier accuracy of *AC+ vs AC−* within the P3b time frame (300-500 ms).

Our second comparison concerned decoding of high vs low uncertainty checkpoints (CP) within the same P3b time frame. The previous finding of a checkpoint-related P3b component as an index of excess information (see above) raised the intriguing question of whether this potential is modulated by global parameters such as context uncertainty. In the absence of adjustment cues within the original paradigm, our initial fMRI study had established differential checkpoint processing as a function of high vs low uncertainty (Kluger & Schubotz, 2017). Consequently, in the present study, we assessed the classifier’s performance decoding high vs low uncertainty checkpoints within the new experimental paradigm. To this end, we sampled regular endings of sequences that did not contain an adjustment cue (REG−), as these events were the precise equivalent of CP from earlier studies. Consolidating previous findings from fMRI and EEG, we thus hypothesised significant above chance classifier accuracy of *CP high vs CP low* within the P3b time frame (300-500 ms).

#### Representational Similarity Analysis

While the decoding analyses described so far allowed us to detect category-specific information in the EEG signal, RSA (Kriegeskorte et al., 2008) provides a framework to test hypotheses about the underlying representational structure of activity patterns. Information carried by a given representation (e.g., of event classes) can be quantified in so-called representational dissimilarity matrices (RDMs). Following the assumption that events with distinct representations are fairly easy to decode (and vice versa), these matrices characterise event representations based on their (dis-) similarities among one another (for review, see Haxby et al., 2014). For hypothesis-based model comparison, multiple hypothetical candidate RDMs can be devised a priori to predict the empirical similarity structure found in the data. By correlating model and neural RDM (done here by means of Pearson correlation), one can then assess the degree to which activity patterns reflect model-inherent structure (in other words, the explanatory power or fit of the respective model RDM).

Two binary model RDMs were set up to predict the empirical representational similarity of event classes in the EEG data (Fig. 3B). While both RDMs commonly modelled third position trials (AC+/−) as highly similar, they differed in their assumptions regarding events at the (potential) endings of a given sequence: The first model RDM (termed *global model*) categorically assumed high similarity within and high dissimilarity between the sampled event classes (AC+/− vs REG+/− vs EXT+/−) and did not account for potential adjustments made in response to incidental adjustment cues. In contrast, the second model RDM - termed *local model* - assumed high similarity of events that violated potentially adjusted predictions following an adjustment cue (REG+, EXT−) in contrast to those that complied with the adjustment cue (REG−, EXT+).

## Results

### Behavioural results

Participants showed an overall high level of performance with a mean P_R_ score of *M_PR_* = .97 (*SD* = .02) during the EEG session, indicating excellent attentiveness throughout the experiment. The repeated-measures ANOVA yielded significant main effects of all three factors on offset latency (*expectation compliance: F*(1, 29) = 53.63, *p* < .001; *adjustment cueing: F*(1, 29) = 19.20, p < .001; *irreducible uncertainty: F*(1, 29) = 4.74, *p* = .003). Neither interaction term reached statistical significance (all *p* > .254). First, replicating behavioural findings from earlier studies (Kluger & Schubotz, 2017; Kluger et al., 2019), a post-hoc t-test revealed significantly quicker button releases after extended (M = 403 ms, SD = 147 ms) than after regular sequences (M = 567 ms, SD = 77 ms; t(29) = −7.32, *p* < .001; factor *expectation compliance)*. Second, significantly shorter offset latencies were found for sequences that contained an adjustment cue (M = 480 ms, SD = 104 ms) than for those that did not (M = 490 ms, SD = 97 ms; *t*(29) = −2.18, *p* = .038; factor *adjustment cueing*). Finally, participants released the response button significantly quicker when irreducible uncertainty was low (M = 473 ms, SD = 98 ms) than when it was high (M = 497 ms, SD = 104 ms; t(29) = −4.38, *p* < .001; factor *irreducible uncertainty*).

Building on analyses of an earlier study (Kluger et al., 2019), we then assessed anticipatory and particularly quick reactions to regular sequence endings as a behavioural marker of adjustment cue recognition. Following event definitions of previous work, these reactions were defined as button releases during the last sequential and the first non-sequential digit (i.e., offset latency ± 500 ms, see Fig. 1C). Since the adjustment cue reliably indicated sequential extensions, participants should respond less frequently to REG+ (i.e. unexpected regular endings) than to REG− sequences (expected regular endings) in this time frame after they successfully learned the adjustment cue. A paired *t*-test supported this hypothesis, revealing significantly fewer responses for REG+ (M = 13.20, SD = 8.40) than for REG− sequences (M = 68.07, SD = 40.25) within the [−500, 500] ms interval (*t*(29) = 9.27, *p* < .001, see Fig. 1).

## Event-related potentials

### UNEXP - EXP

Building on previous EEG results (Kluger et al., 2019), we first aimed to replicate findings of an N400 component for the direct comparison of unexpected and expected outcomes. Accordingly, we included all time points between 300 and 500 ms in a one-sided whole-brain analyses (6262 comparisons in total). As hypothesised, unexpected outcomes (EXT−, REG+) were found to elicit a significant N400 over parieto-central electrodes peaking around 442 ms (Fig. 2).

**Figure 2.**
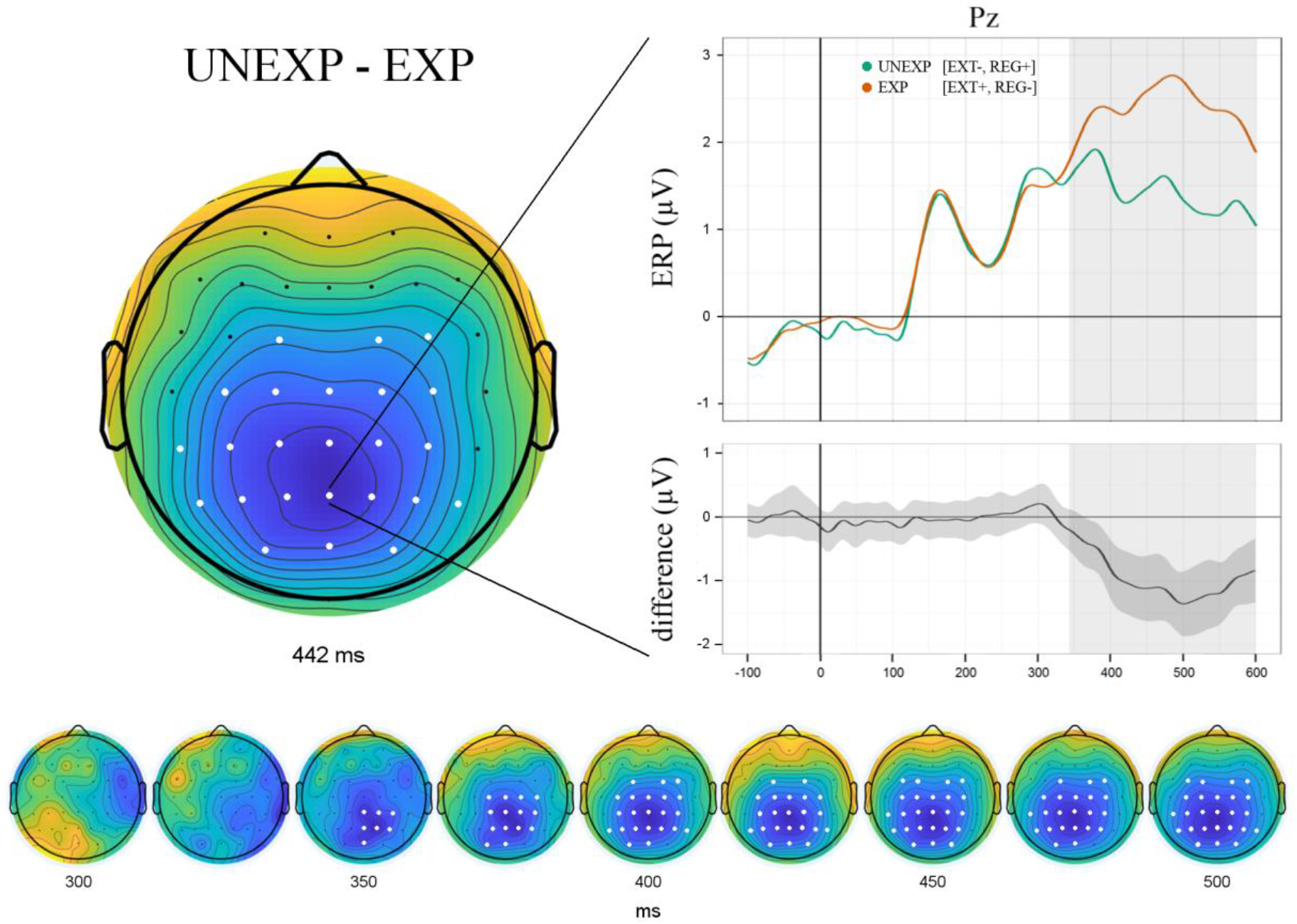
Direct comparison of unexpected (EXT−, REG+) and expected outcomes (EXT+, REG−) revealed a significant N400 component peaking around 442 ms over parieto-central electrodes. The difference curve includes M ± STD of individual data; bottom panel shows component evolution over time.

## Multivariate classification

### AC+ vs AC−

In a first classification analysis, we trained the classifier to decode third-position events that contained an adjustment cue (AC+) from those that did not (AC−, see Fig. 1B). Group-level classification analysis revealed temporal clusters with significant above chance decoding accuracy between 140 - 310 ms as well as between 330 – 420 ms (Fig. 3A). Intriguingly, the second cluster particularly supported our hypothesis regarding the P3b time frame for this comparison. In addition, two brief significant clusters were found from 570-590 ms and from 770-800 ms, respectively. However, since these time frames already fell within the presentation of the next stimulus, these results should be interpreted with caution.

**Figure 3.**
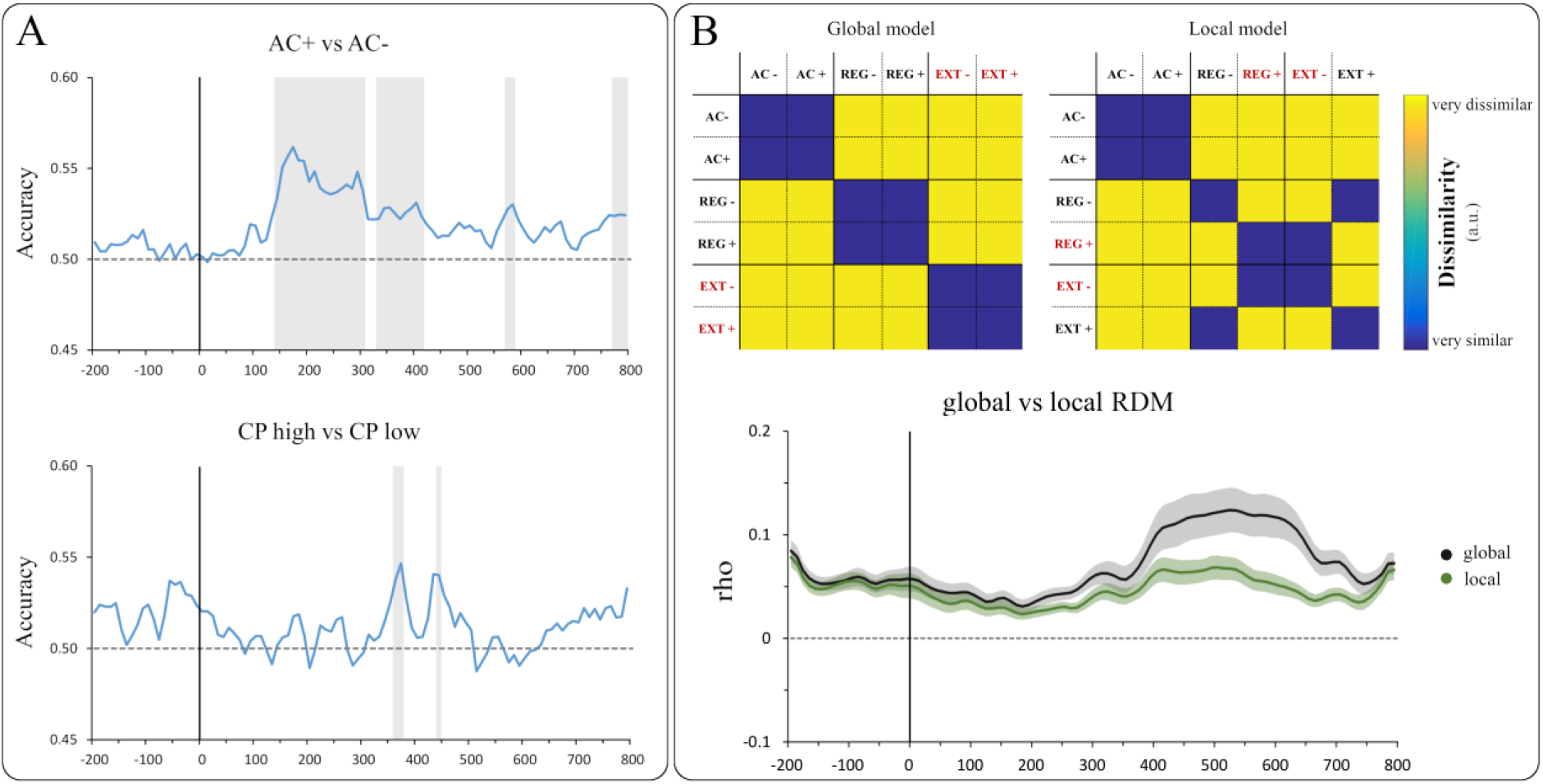
(A) Group-level decoding accuracies over time. Shaded areas indicate significant temporal clusters. (B) Top: Model RDMs predicting representational similarity in the EEG data based on distinct event characteristics. Globally (left) and locally (right) unexpected events are marked in red, respectively. Bottom: Group-level correlations of both model RDMs with the neural RDM. Shaded area around the curve shows SEM of group average correlation. AC = adjustment cues, CP = checkpoints, REG = regular endings, EXT = sequential extensions.

### CP high vs CP low

Extending the narrative of previous studies (see Kluger & Schubotz, 2017) on modulation by context statistics, the second classification analysis concerned checkpoints (CP) under high and low uncertainty, respectively. As the present experimental design featured locally informative adjustment cues, regular sequence endings with no prior adjustment cue (REG−) immediately corresponded to CP events sampled in previous studies (Kluger & Schubotz, 2017; Kluger et al., 2019). Accordingly, we trained the classifier to decode these CP in high uncertainty contexts from those in low uncertainty contexts. As hypothesised, group-level classification analysis revealed significant temporal clusters within the P3b time frame (360-380 ms). A second small cluster was found for the time frame between 440-450 ms (Fig. 3A).

### Representational Similarity Analysis

We employed searchlight RSA to compare two binary representational dissimilarity matrices (RDMs) modelling representational similarity of event classes in the EEG data. Both the global and the local model were found to be significantly correlated with the neural RDM over the entire time course of [−200, 800] ms (Fig. 3B). Direct model comparison by means of a Wilcoxon signed-ranks test revealed that the global model showed an overall better model fit to the EEG data (mdn = 0.06) than the local model (mdn = 0.05; *Z* = 8.86, *p* <.001), particularly apparent later in the time series (around 400-700 ms). In line with central propositions of predictive processing accounts, the statistical comparison of both model RDMs demonstrates that the global event structure acquired through statistical learning took precedence over online information conveyed by intermittent adjustment cues. In other words, statistical learning of event categories (REG, EXT) was not cancelled out by temporary shifts in expectations but remained the strongest predictor of overall similarity in EEG activation patterns.

## Discussion

Predictive brain processing vitally relies on top-down hierarchical models continuously matched against incoming sensory information. In case of deficient model assumptions, mismatch signals termed prediction errors (PE) are propagated upstream to instigate model updating for future occasions. Recent fMRI and EEG studies (Kluger & Schubotz, 2017; Kluger et al., 2019) have demonstrated exploitation of information from model-compliant events, particularly in contexts of high uncertainty. Building on previous work highlighting such global modulations in predictive processing, the present study was conducted to assess the influence of locally available information. Therefore, we introduced adjustment cues which interfered with the learned statistical task structure, effectively calling for a flexible on-line adjustment of expectations.

Multivariate classification analysis revealed that adjustment cues could be reliably decoded from uninformative events at the same position within the P3b time frame. Moreover, events that were unexpected globally (according to task structure) or locally (as indicated by adjustment cues) elicited a mismatch-related N400 component when compared to expected events. Combined with behavioural data, these findings demonstrate that local cueing information was efficiently exploited to adapt to current task requirements.

Aside from on-line adjustments prompted by local cues, global uncertainty was found to modulate the functional signature of checkpoints: Consolidating previous evidence from fMRI (Kluger & Schubotz, 2017) and EEG (Kluger et al., 2019), checkpoints were distinctly processed under high (vs low) uncertainty within the P3b time frame, as indicated by significant multivariate classification accuracy. Finally, representational similarity analysis (RSA) was used to compare two dissimilarity models representing global and local contingencies, respectively. While both models were found to be significantly correlated with the neural dissimilarity matrix, the global model was found to fit the neural data better at a later time frame (400 – 700 ms). Conceivably, higher cognitive operations reflected in this time frame are more likely to incorporate global information acquired through long-term statistical learning.

### Local information prompts on-line adjustments

It was one of the main aims of this study to assess the influence of two separate sources of information in predictive processing. First, in the present experimental context, global task structure and transition probabilities are acquired through statistical learning, starting with the training session. Higher-order information also comprised different levels of irreducible uncertainty, that is, how frequently implicit expectations regarding sequence length (“… usually five digits long”) were violated by sequential extensions. Secondly, *local* information was introduced in the form of adjustment cues reliably indicating a specific outcome (i.e., an extension of that sequence). This information thus modified the statistically learned contingencies and called for a rapid adjustment of expectation in an ongoing sequence.

Effective exploitation of adjustment cues was reflected in behavioural response patterns at regular sequence ends: Participants responded less frequently to unexpected regular endings (REG+) within a time frame of ± 500 (compared to regular endings, REG−). This time frame was used in a previous study to mark anticipatory and particularly quick reactions, indicating behavioural facilitation effects of cue recognition (Kluger et al., 2019). In the present study, the preceding adjustment cue had reliably indicated an extension, rendering the overall more probable regular ending locally improbable. Recognising adjustment cue information, participants were conceivably less confident to prepare an anticipatory or quick release of the response button. This finding fits well with previous results consistently showing response slowing for sequences that - much like present REG+ trials - ended earlier than indicated by the preceding cue (Kluger & Schubotz, 2017; Kluger et al., 2019).

On the neural level, two key findings provided critical insight into the role of adjustment cue processing: First, supporting our hypothesis, ERP analysis revealed a significant N400 component for locally unexpected (vs expected) sequential events. Critically, unexpected events comprised REG+ and EXT− trials, thus accounting for adjusted expectations due to cue information1. This finding replicates results from an earlier study (Kluger et al., 2019) in which we established N400 as the proper mismatch correlate in sequential prediction. The N400 component has been reported to be elicited by a multitude of sensory events and varies in amplitude as a function of event-bound surprise in language (Rabovsky et al., 2018; Szewczyk & Schriefers, 2018) and arithmetic problems (Galfano et al., 2004; Niedeggen et al., 1999). More generally, it is discussed as a modality-independent index of conceptual representations which need revision in light of violations (for review, see Kutas & Federmeier, 2011). In sum, our interpretation of N400 as a correlate of locally induced mismatches further corroborates the component’s role in integrating incongruous information.

Second, as hypothesised, multivariate pattern classification revealed significant decoding of AC+ and AC− trials within the P3b time frame. P3b has consistently been reported for informative, task-relevant stimulus evaluation (Huang et al., 2015; Katayama & Polich, 1998). Critically, our findings emphasise that stimuli do not necessarily have to violate model assumptions to be informative. Previous studies have shown P3b to be evoked by an informative sequence of standard trials acting as a ‘secondary target’ (Fogelson et al., 2009) and even in the absence of stimuli if the omission was informative (Nieuwenhuis et al., 2011). Functionally, P3b appears to facilitate adequate responses following behaviourally relevant stimuli (Verleger et al., 2005). In the present study, information gained from adjustment cues modified global contingencies and facilitated preparation of the adequate response (i.e., most probably keeping the finger on the response button for an extended sequence).

Adjustment cues (AC+) thus bear a striking conceptual resemblance to checkpoints (CP) defined in previous studies: In both cases, model-compliant sensory input of probabilistic events can be exploited to evaluate the predictive model in the absence of a prediction error. Critically, as reflected by significant decoding within the P3b time frame, this was not the case for uninformative third-trial events (AC−). Sensory input at these positions was model-compliant but did not provide excess local information, making AC− trials comparable to non-informative standard trials.

### Global effects of contextual uncertainty

To assess the effect of irreducible uncertainty on checkpoint processing, we once again employed multivariate pattern classification for decoding of CPs under high and low uncertainty, respectively. In order to increase comparability between studies, we restricted the present analysis to checkpoints as they were defined in earlier work, namely regular endings of sequences that did not contain an adjustment cue (REG−). Supporting our hypothesis, we found clusters of significant decoding accuracy within the P3b time frame. This finding considerably adds to earlier results from two directions: First, as mentioned above, our previous EEG study revealed P3b as the correlate of excess information gain at checkpoints, effectively separating it from non-informative analogues (Kluger et al., 2019). Second, a previous fMRI study had established a functional network exhibiting increased neural activity for CP processing under high (vs low) irreducible uncertainty (Kluger & Schubotz, 2017). Notably, this network prominently included the temporo-parietal junction (TPJ), a conceivable cortical source of P3b (Donchin & Coles, 1988). Altogether, the present finding suggests a modulatory influence of contextual uncertainty on information exploitation at checkpoints which translates to distinct correlates within the P3b time frame: As high uncertainty blocks were defined by a higher proportion of extended sequences, validity of the initial model (“… usually five digits long”) was reduced in these blocks. Therefore, sites of supposed sequence endings were particularly informative whether a motor response was in fact required (REG−) or not (EXT−). In line with accounts of contextual updating ascribed to both P3b (Kimura et al., 2010; Verleger et al., 2016) and its suggested source in TPJ (Geng & Vossel, 2013), we thus interpret the present uncertainty effect as a correlate of excess information being exploited to estimate global context features and adapt behaviour accordingly.

It is worth noting that, since AC+ trials reliably indicated extensions, high uncertainty blocks necessarily contained a higher absolute number of adjustment cues than low uncertainty blocks to ensure constant cue validity (see Fig. 1B). In other words, whereas the global portion of extended sequences was higher in these blocks, local probabilities after the fifth sequential position remained unchanged. The finding of significant checkpoint decoding under high vs low uncertainty thus suggests that local information did not fully cancel out global contingencies and that the role of CP as informative reference points should be even more evident in contexts without local adjustment cues (see Limitations and future directions).

### Predictive efficiency across time scales

Representational similarity analysis of a local and a global model RDM revealed significant correlation with neural data for both models. Pre-eminence of the global model (apparent especially in a later time frame around 400 - 700 ms), however, leaves room for discussion and motivates further research. Critically, the two models differed with regard to which events were jointly classified as unexpected: The local model reflected adjusted probabilities as a function of on-line cues, grouping REG+ and EXT− occurrences as locally unexpected events. In contrast, the global model did not account for any adjustments but rather modelled all extensions (EXT+, EXT−) as similar and as distinct from regular endings (REG+, REG−). The differential modelling of unexpectedness in the model RDMs as well as the late timing of the difference in model fit strongly suggest these RSA results to be related to the N400 finding reported above: While locally unexpected events could be reliably decoded from expected events in the preceding classification analysis, the higher fit of the global RDM shows that - across all event classes - late potentials reflected an even higher (dis-)similarity of globally unexpected (vs expected) events. In other words, whereas local information was undoubtedly exploited for on-line adjustments in prediction and behaviour, it did not fully overwrite statistically learned global contingencies. Previous studies have highlighted this multi-level hierarchical organisation by demonstrating distinct error signals for the violation of local probabilities and global rules (Bekinschtein et al., 2009; Wacongne et al., 2011), effectively showing the distinction between unexpected and expected mismatch signals. Consequently, present RSA results corroborate a fundamental notion of hierarchical predictions, namely that global reference frames always rank higher than local expectations in predictive processing. Fittingly, data from previous N400 studies revealed that when information from two different levels (such as word- and sentence level) is available, higher-level context effects tend to take precedence over lower-level ones (Kutas, 1993; van Petten, 1993). Indeed, frequently re-learning by overriding global probabilities seems quite costly in terms of computational efficiency. Instead, we may establish a global situational model (in this case reflecting the expectable sequence length) which can be locally modified - but not overturned - in the light of additional information. In line with interpretations of N400 as an index of information integration (which is more effortful for incongruous input, see above), our findings overall indicate that the effort of integration is invested in a model of global contingencies rather than representing local fluctuations.

Underscoring the precedence of global over local information, successful multivariate decoding of high vs low uncertainty checkpoints was particularly instructive in that local information from adjustment cues could have indeed cancelled out the functional significance of checkpoints altogether: Extensions were in fact more frequent under high (vs low) uncertainty, but were still reliably preceded by adjustment cues. Therefore, when no adjustment cue was presented, local probabilities of extensions were factually identical in high and low uncertainty contexts (see Fig. 1B). Distinct behavioural and functional indices of adjustment cue information notwithstanding, differential processing of high vs low uncertainty checkpoints thus demonstrates how global models are - to a certain extent - shielded against energetically costly refinement.

### Limitations and future directions

The aim of the present study was to investigate the interplay of local and global information in predictive processing. To this end, we employed local adjustment cues which were incidentally presented to interfere with statistically learned, global contingencies. In order to maximise frequency for the events of interest, we were not able to analyse sequential standard trials due to their close temporal proximity to other sampled events. However, as the present study has demonstrated differential processing of informative vs non-informative cue positions, it would be revealing to assess the extent to which these positions are processed differently from non-informative sequential standards: Even if no cue is presented, the *potential* information gain at respective sequence positions may be sufficient to single out these events compared to deterministic standard trials. This kind of functional prominence, in turn, conceivably depends on how vital potential information from cue positions would be in the current task context. Conceivably, the more instructive adjustment cues are, the more resources will presumably be allocated to sequential positions from which such information could potentially be gained, irrespective of the actual outcome. To this end, future studies should assess neural correlates not only in response to, but also leading up to, these events. Upcoming efforts should assess expectations prior to potentially informative sequential positions by means of pre-stimulus ERPs or time frequency analyses. One promising candidate marker of information expectation is the stimulus-preceding negativity (SPN; see Mnatsakanian & Tarkka, 2002), a slow potential shown to increase in amplitude prior to the presentation of particularly informative events (Morís et al., 2013). It remains to be examined whether potential cue positions are preceded by correlates of increased expectations when compared to non-informative standard trials. In a similar vein, as distinct components of predictive processing have been suggested to be transmitted by separate oscillatory rhythms (e.g., Arnal & Giraud, 2012), assessment of pre-stimulus differences in the power of β- and γ-frequency bands would considerably add to the understanding of adjustment cue and checkpoint processing.

### Conclusion

Model-compliant events have previously been established as critical reference points informing predictive processing in the absence of prediction errors. A similar functional role on the local level is now suggested for a different class of model-compliant events: Accounting for information from local *adjustment cues*, we were able to replicate a significant N400 component as an index of mismatch-induced model adaptation. Furthermore, using multivariate pattern classification, sequential positions providing excess local information could be reliably decoded from those that did not within the P3b time frame. Global effects of irreducible uncertainty on checkpoint processing remained, as reflected by significant classification of high vs low uncertainty checkpoints in the same time frame. Representational similarity of a global model was found to fit the neural data better than a local model, suggesting that adjustments reflected by late ERPs (e.g., N400) are made to a higher-level model of global contingencies. Overall, our pattern of results suggests that just like error-induced model adaptation, model evaluation is not limited to either local or global information. Instead, the overall hierarchical organisation of predictive processing implies that model evaluation can occur at different levels of the processing hierarchy. Intriguing research questions remain with regard to the precise interplay of local vs global information sampling.

## Acknowledgements

We would like to thank Monika Mertens, Katharina Thiel, Corinna Gietmann, and Marie Kleinbielen for their help during data collection as well as Joachim Gross for insightful comments on an earlier draft of the manuscript.

## Footnotes

1 Note that expected (REG−, EXT+) and unexpected (REG+, EXT−) event classes were defined by means of a weighted average of trials, precluding ERP effects to be caused by mere differences in trial numbers.

